# Measuring Granger-causal effects in multivariate time series by system editing

**DOI:** 10.1101/504068

**Authors:** Roberto D. Pascual-Marqui, Rolando J. Biscay, Jorge Bosch-Bayard, Pascal Faber, Toshihiko Kinoshita, Kieko Kochi, Patricia Milz, Keiichiro Nishida, Masafumi Yoshimura

**Affiliations:** The KEY Institute for Brain-Mind Research, Department of Psychiatry, Psychotherapy, and Psychosomatics, University Hospital of Psychiatry, Zurich, Switzerland; Department of Neuropsychiatry, Kansai Medical University, Osaka, Japan; Centro de Investigacion en Matematicas, Guanajuato, Mexico; Departamento de Neurobiologia Conductual y Cognitiva, Instituto de Neurobiologia, Universidad Nacional Autonoma de Mexico, Queretaro, Mexico

## Abstract

What is the role of each node in a system of many interconnected nodes? This can be quantified by comparing the dynamics of the nodes in the intact system, with their modified dynamics in the edited system, where one node is deleted. In detail, the spectra are calculated from a causal multivariate autoregressive model for the intact system. Next, without re-estimation, one node is deleted from the model and the modified spectra at all other nodes are re-calculated. The change in spectra from the edited system to the intact system quantifies the role of the deleted node, giving a measure of its Granger-causal effects (CFX) on the system. A generalization of this novel measure is available for networks (i.e. for groups of nodes), which quantifies the role of each network in a system of many networks. For the sake of reproducible research, program codes (PASCAL), executable file, and toy data in human readable format are included in the supplementary material.

## 2. Introduction

Research in human brain function, based on neuroimaging techniques that yield distributed signals of cortical activity, is mostly dedicated to function localization and to connectivity.

These two aspects have for a long time remained as separate goals in research, as can be appreciated from the following two quotations:

1. Friston 2011: “Imaging neuroscience has firmly established functional segregation as a principle of brain organization in humans. The integration of segregated areas has proven more difficult to assess.”
2. Seth et al 2015: “A key challenge in neuroscience and, in particular, neuroimaging, is to move beyond identification of regional activations toward the characterization of functional circuits underpinning perception, cognition, behavior, and consciousness.”

In this paper, a novel method of analysis is presented that attempts to bring together in a unified manner the influence that any one cortical region has on all other cortical regions, as mediated by the causal effective connectivity pattern of the system.

Consider a system consisting of *p* ≥ 2 nodes, where time series measurements are available at each node. The main example of interest here consists of high time resolution signals of electric neuronal activity at *p* ≥ 2 cortical regions.

In this context of brain activity signals, under wide sense stationarity (Brillinger 2001), the power spectrum at each node provides localized functional information, while the coherencies provide information on *functional connectivity*. One of the most commonly used estimators for these measures is the non-parametric cross-spectrum obtained from the average periodogram.

On the other hand, information on *effective causal connectivity*, in the sense of Granger, can be derived from the parameters of a multivariate autoregressive (MAR) model (Granger 1969; Geweke 1982, 1984). As can be seen from the published literature (see e.g. Valdes et al 2011; Sameshima and Baccala 2014; Seth et al 2015), the MAR model is widely used, but almost exclusively for connectivity inference, and not for the characterization of localized function, even though the MAR model also provides an estimator for the power spectra.

In the novel definition introduced here, the spectra are calculated from a MAR model for the intact system of “*p*” nodes. Next, one node is deleted from the model, and without re-estimation of the MAR parameters, the modified spectra at all other nodes are re-calculated. The change in spectra from the edited system to the intact system quantifies the role of the deleted node, giving a measure of its causal effects (CFX) on the system.

## 3. The multivariate autoregressive (MAR) model and Granger-causality in the time domain

A stable multivariate autoregressive model of order *q* ≥ 1, for *p* ≥ 2 time series 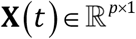, is written as:

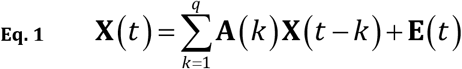

where “*t*” denotes discrete time, 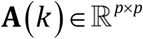 are the causal autoregressive coefficients, and 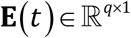 is the innovations (noise) vector with covariance matrix 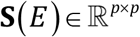. The autoregressive coefficients *A*_*ij*_(*k*) i.e. the element (*i, j*) of the matrices **A**(*k*), quantify the direct causal influence *j* → *i*, i.e. from node “j” to node “i”, for *i* ≠ *j*. Note that in practice, a simple least squares fit can be used to estimate the parameters **A**(*k*) and **S**(*E*) in Eq. 1.

If *A*_*ij*_(*k*) ≠ 0, for *i* ≠ *j*, for any lag “*k*”, then “j” causes “i” in the sense of Granger (Granger 1969; Lütkepohl 2005).

## 4. The MAR model and its Granger-causality in the frequency domain

The frequency domain representation corresponding to Eq. 1 is:

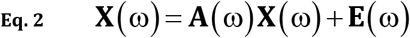

where “ω” denotes discrete frequency, and 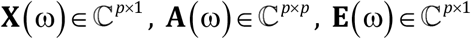 are the respective discrete Fourier transforms.

In the frequency domain, if *A*_*ij*_(ω) ≠ 0, for *i* ≠ *j*, then “j” causes “i” in the sense of Granger at frequency ω (see e.g. Schelter et al 2009; Pascual-Marqui et al 2014).

## 5. Power spectra mediated by causal connectivities

From Eq. 2, the Hermitian covariance for 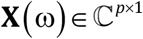, i.e. its cross-spectral matrix, is:

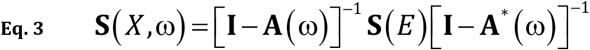

where the superscript “*” denotes matrix transpose and complex conjugate, the superscript “-1” denotes matrix inversion, and **I** is the identity matrix.

The real-valued diagonal elements of **S**(*X*, ω), i.e. *S*_*ii*_(*X*, ω) for *i* = 1…*p*, correspond to the spectral power for the i-th node.

Note that the spectral power at any particular node, at any frequency, is a non-linear function of the set of all the causal connectivity coefficients in the matrix **A**(ω).

## 6. Causal effects (CFX) by system editing

The next step in quantifying the causal effect of the j-th node on all other nodes consists of deleting the j-th node from the system, without re-estimation of the system parameters.

Deletion of the j-th node is implemented by setting the j-th row and column to zero in the MAR parameter matrices and is denoted as **A**_〈*j*〉_ and **S**_〈*j*〉_.

Thus, the power spectra for the remaining nodes in the edited system without the j-th node correspond to the diagonal elements (excluding the j-th diagonal element) of the matrix:

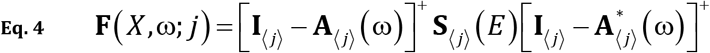

where the superscript “+” denotes the Moore-Penrose pseudoinverse.

Finally, “Granger-causal effects by system editing”, denoted as CFX, is defined as the (causal) change of power spectra from the system without the j-th node, to the system with the j-th node:

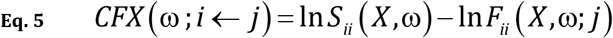

for all nodes *i* = 1…*p* such that *i* ≠ *j*.

*CFX*(ω; *i* ← *j*) in Eq. 5 quantifies the causal effect of node “j” on the observed activity at node “i”.

Note that this measure can be positive or negative. This means that the addition of a deleted node can causally increase or decrease the spectral power at the other nodes, differently for each frequency.

Also note that one useful property of the difference of logarithms in Eq. 5 is its invariance to scale, which makes it dimensionless, allowing a comparison of effects on all nodes independent of scale.

## 7. CFX Statistics

There are typically two levels of statistical tests in the field of neuroimaging, applied to both localization data and to connectivity data.

First level (or single subject) statistics applies to data from a single subject, corresponding to, e.g., time series measurements in a single subject. Knowledge of the statistical distribution of the collection of estimated values of causal effects CFX (Eq. 5) for all pairs of nodes (*i, j* = 1…*p, i* ≠ *j*) and for all discrete frequencies ω, is required. Unfortunately, this is not readily available.

However, note that the CFX measure in Eq. 5 is the logarithm of a “ratio of means”. This is because the spectral density is equivalent to statistical variance, which is at the same time the “mean value” of the squared modulus of a complex valued Fourier transform coefficient.

The logarithm of a “ratio of means” is a typical statistic for which the effect size has been well studied, see e.g. Hartung et al 2008, page 110 and 111 therein. It is shown there that:

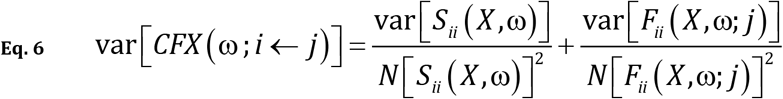

where “var” is the variance operator, and “N” is the sample size. Under the assumption of a Gaussian distribution, the variance of the variance, i.e. the variance of the spectral density is known:

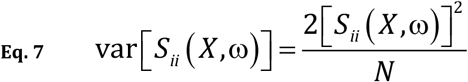

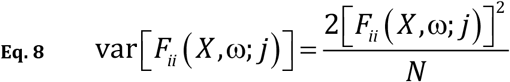

Plugging Eq. 7 and Eq. 8 into Eq. 6 simplifies to:

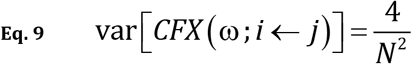

with standard deviation:

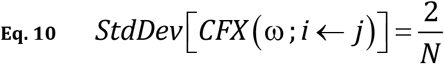

From Eq. 5 and Eq. 10, a t-statistic has the form:

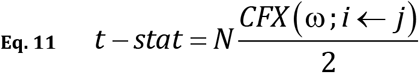

However, the effect size, which is independent of sample size, expressed in the form of “Cohen’s d”, is preferred in this case:

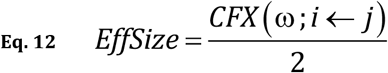

See e.g. Cohen 1988, and more recently Poldrack et al 2017.

The well know “Cohen’s d” thresholds for effect size in this case are:

Small effect size: 0.2
Medium effect size: 0.5
Large effect size: 0.8

This gives the following simple rules of thumb for the size of CFX:

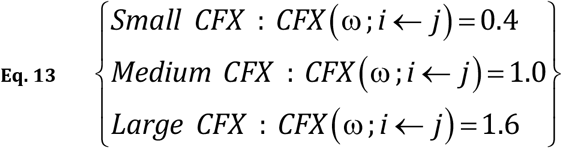

The rules of thumb in Eq. 13 can be applied to first level, single subject analyses.

Now consider the case of second level statistics, i.e. group data. The simplest situation corresponds to a single group composed of “M” subjects, for which estimated values of causal effects CFX (Eq. 5) are available for all subjects (*k* = 1…*M*), for all pairs of nodes (*i* = 1…*p, i* ≠ *j*), and for all discrete frequencies ω, denoted as: *CFX*_*k*_(ω; *i* ← *j*).

For the given sample of size “M”, tests for zero mean, with null hypotheses:

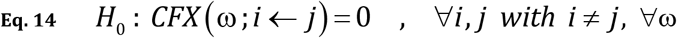

can be performed, using, for instance, simple t-statistics. Finally, correction for multiple testing can be achieved by non-parametric randomization of the maximum t-statistic, which in addition does not require the assumption of a Gaussian distribution (see e.g. Nichols and Holmes 2002).

The generalization to more complicated forms of second level statistics, for instance, with several independent groups, is straightforward (see e.g. Nichols and Holmes 2002).

## 8. A toy example

In this section, use will be made of the same toy example that was designed and studied in Pascual-Marqui et al 2014.

The toy example corresponds to five signals generated from a stable second order MAR model:

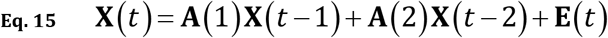

with 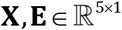, 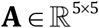, with zero mean innovations having a covariance matrix equal to the identity matrix:

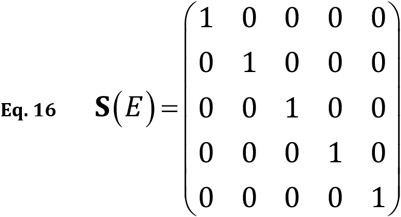

The causal autoregressive coefficients used in the Pascual-Marqui et al 2014 study were:

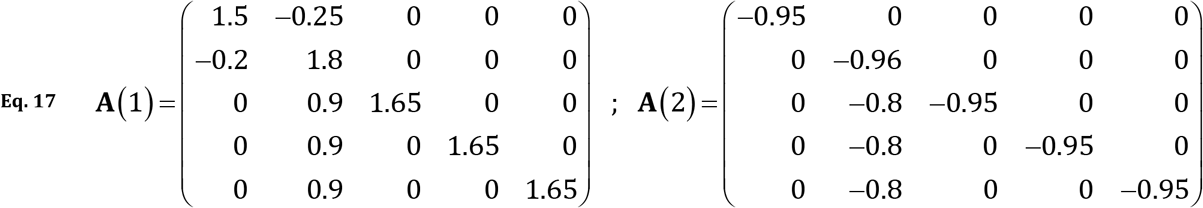

The connections in this system, which are schematically shown in Figure 1, are as follows:

- Node 1 sends to node 2,
- Node 2 sends to nodes 1, 3, 4, and 5

**Figure 1:**
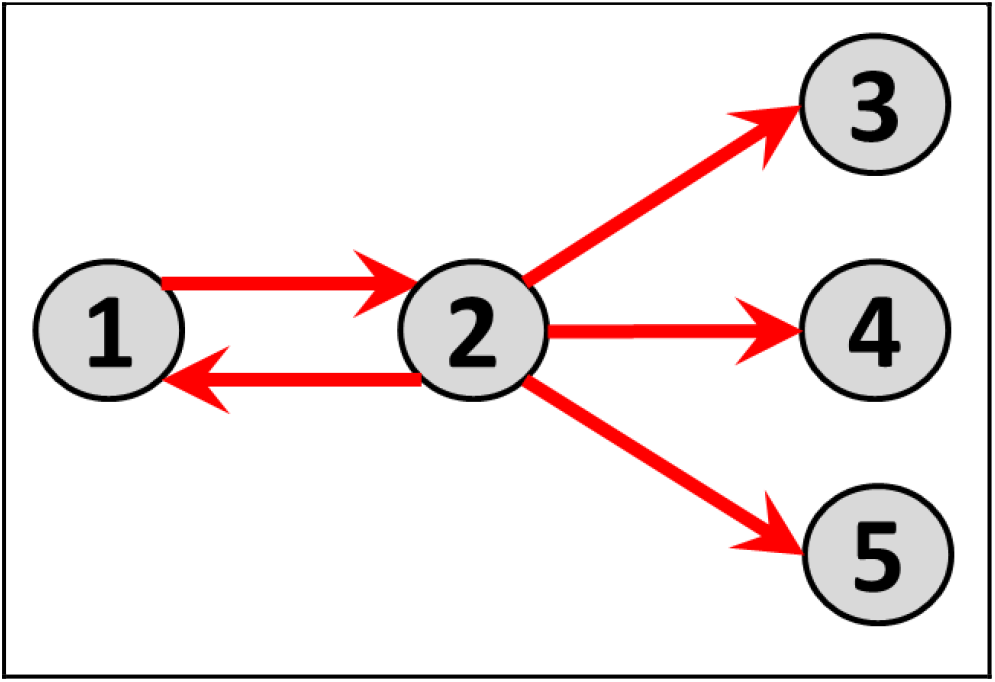
Schematic outline of the causal connections in the toy example.

Eq. 15, Eq. 16, and Eq. 17 were used to generate a realization of the five signals, consisting of 25600 time samples at a nominal sampling rate of 256 Hz. Figure 2 shows a 4 second segment of the five signals.

**Figure 2:**
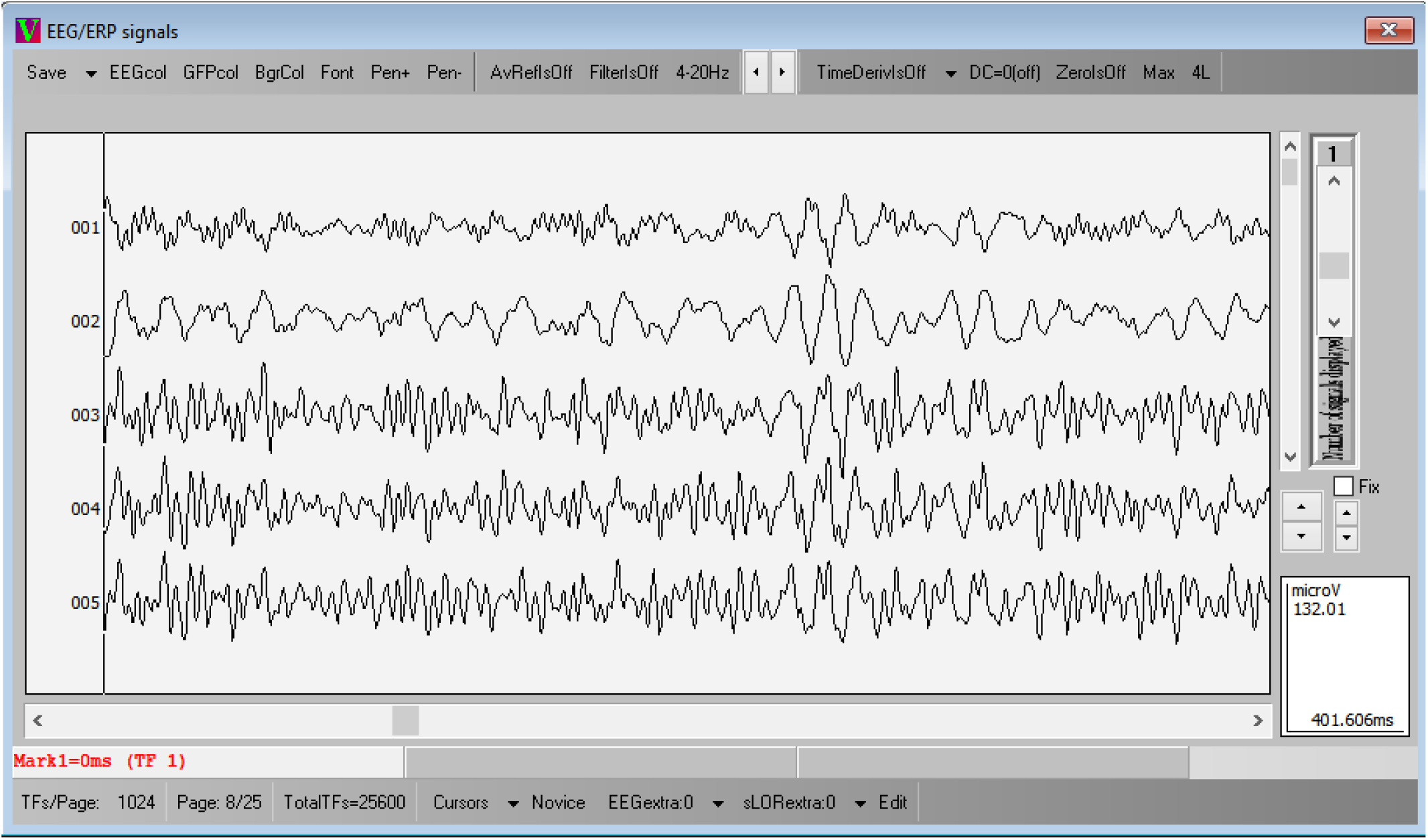
Display of four seconds (1024 time samples) of the five signals as generated by Eq. 15 and Eq. 17.

Figure 3 **(A)** shows the log-power spectra corresponding to the data generated by the MAR model (plugging Eq. 16 and Eq. 17 into Eq. 3). The spectral peaks are distributed as follows: (X_1_ and X_2_ at 8 and 32 Hz), and (X_3_, X_4_, X_5_ at 8, 23, and 32 Hz).

**Figure 3:**
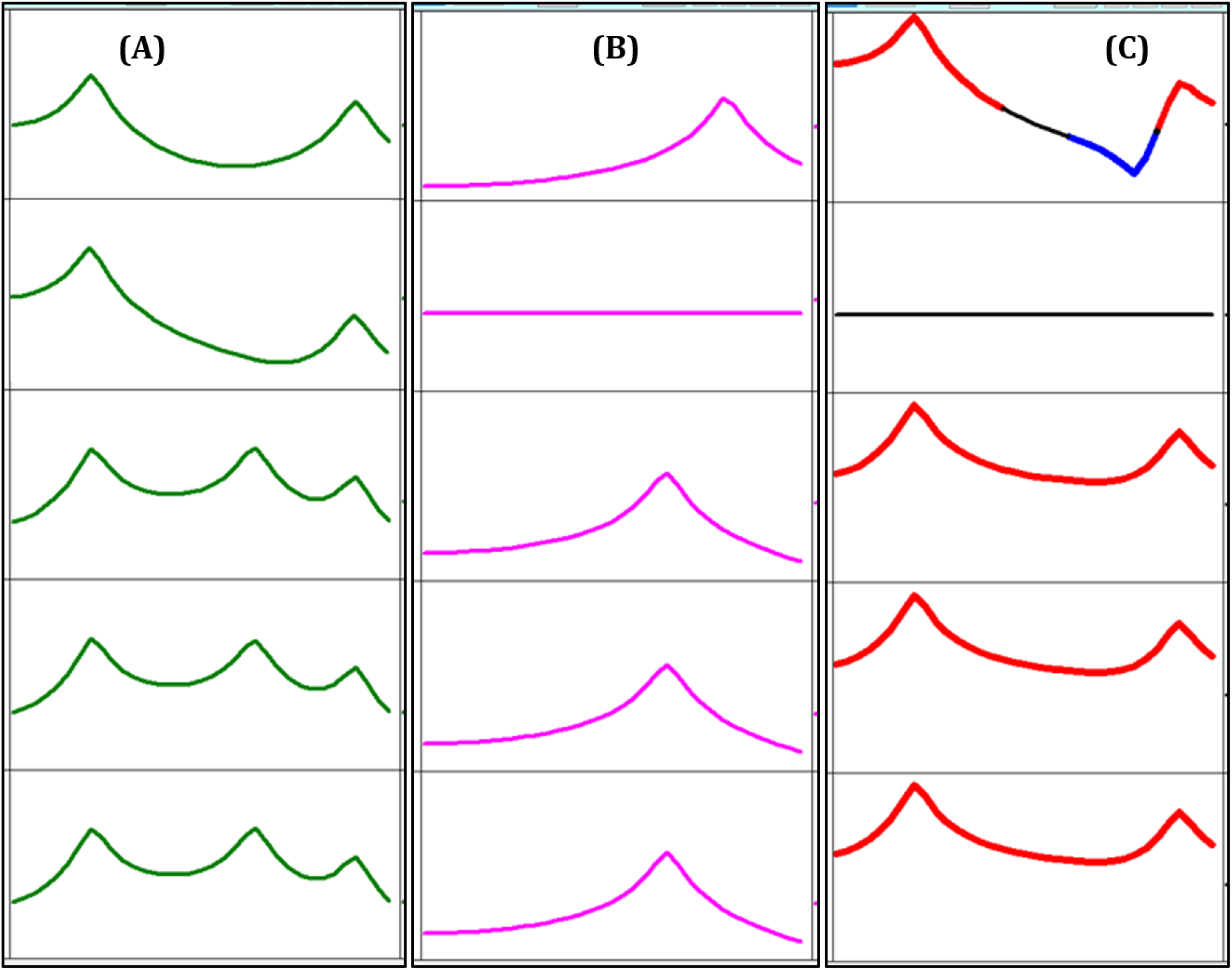
**(A)** Log-power spectra for the intact, non-edited system, obtained from the MAR model (plugging Eq. 16 and Eq. 17 into Eq. 3). The spectral peaks are distributed as follows: (X_1_ and X_2_ at 8 and 32 Hz), and (X_3_, X_4_, X_5_ at 8, 23, and 32 Hz). **(B)** Log-power spectra for the edited system, with node 1 deleted, obtained from the MAR model obtained by plugging Eq. 18 and Eq. 19 into Eq. 4. The spectral peaks are now distributed as follows: (X_2_ at 17 Hz), and (X_3_, X_4_, X_5_ at 17 and 23 Hz). Finally, the difference of log-spectra of the intact system minus the edited system is displayed in **(C)**, which shows that the role of node 2 in the system consists of activating 8 Hz and 32 Hz oscillations at nodes 1, 3, 4, and 5, and also consists of deactivating 17 Hz oscillations at node 1. In column **(C)**, red color corresponds to positive values of CFX, and blue color to negative values of CFX. The horizontal frequency axis spans 1 to 35 Hz.

In the next step, as an interesting example, node 2 is deleted from the system, which corresponds to the following MAR model parameters:

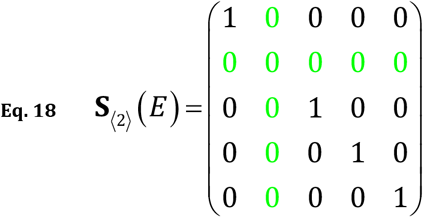

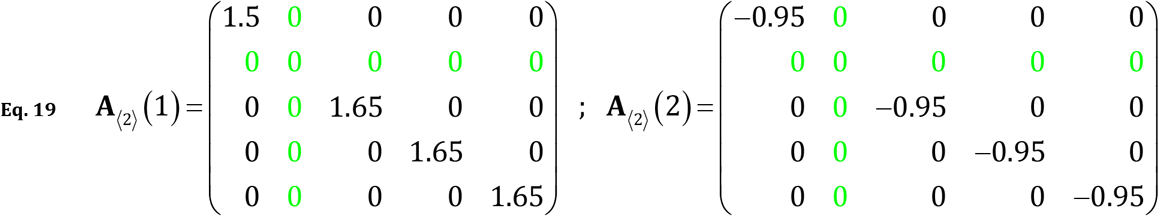

Plugging Eq. 18 and Eq. 19 into Eq. 4 gives the log-power spectra for nodes 2, 3, 4, and 5 in the edited system, which is shown in Figure 3 **(B)**, where spectral peaks are now distributed as follows: (X_1_ at 8 Hz), and (X_3_, X_4_, X_5_ at 23 Hz). Finally, the difference of log-spectra of the intact system minus the edited system is displayed in Figure 3 **(C)**, which shows that the role of node 2 in the system consists of activating 8 Hz and 32 Hz oscillations at nodes 1, 3, 4, and 5, and also consists of deactivating 28 Hz oscillations at node 1. In Figure 3 **(C)**, red color corresponds to positive values of CFX, and blue color to negative values of CFX.

The complete analysis, where each node, one-by-one, is deleted from the system, is shown in Figure 4 and Figure 5.

**Figure 4:**
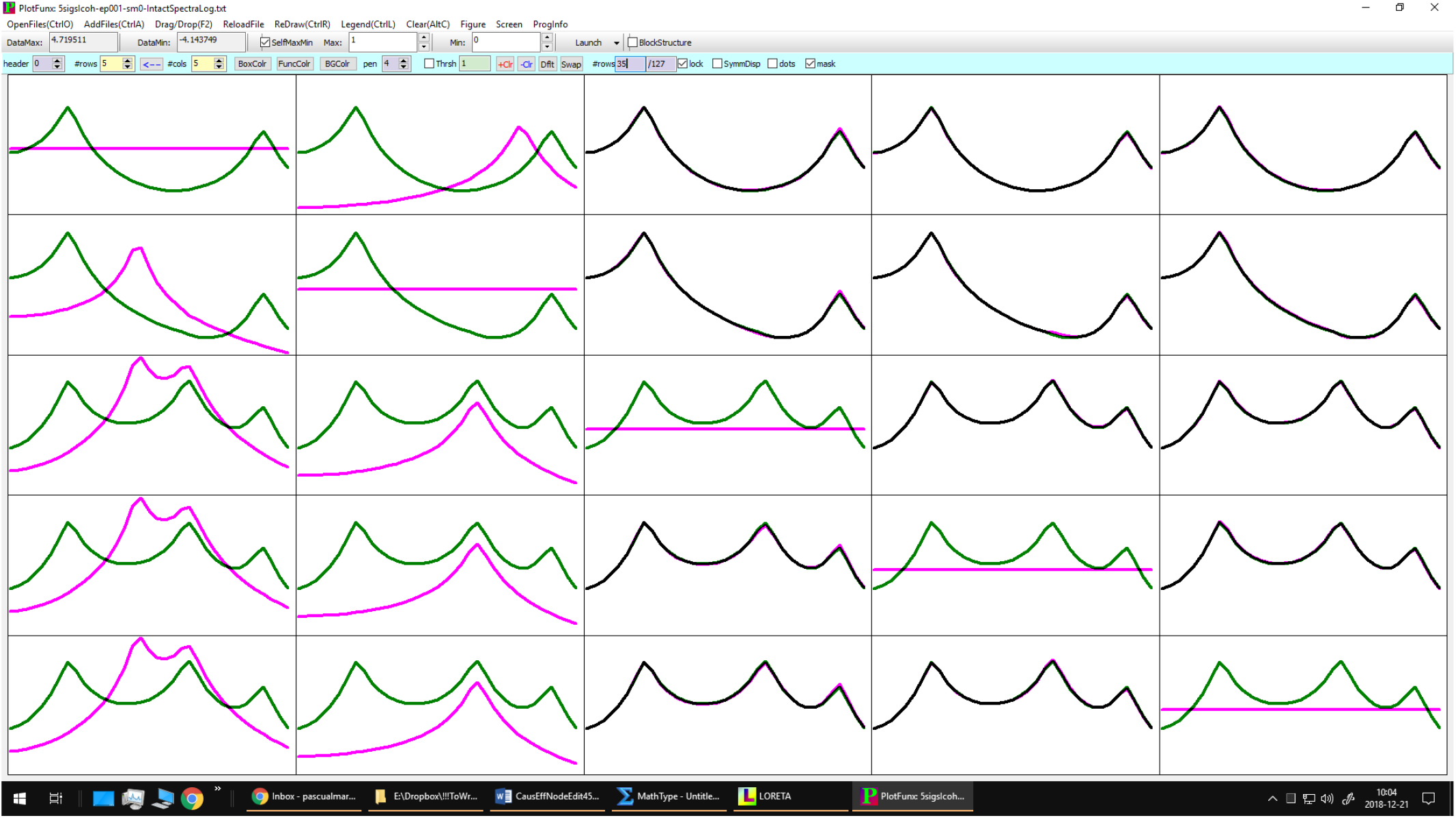
A matrix of curves of size 5×5 is displayed. The magenta colored curve in the i-th row and j-th column corresponds to the power spectra at the i-th node in the edited system where the j-th node has been deleted (obtained from Eq. 4). The green colored curve in the i-th row corresponds to the power spectra at the i-th node in the intact system, obtained from Eq. 3. Note that all green curves for the intact system in the i-th row are identical, independent of the column location. Curves in color black correspond to overlap of the two spectra for the edited and intact systems. The curves from column 2 appear in Figure 3 (A) and (B). The horizontal frequency axis spans 1 to 35 Hz. The vertical axis spans −4.1 to +4.7.

**Figure 5:**
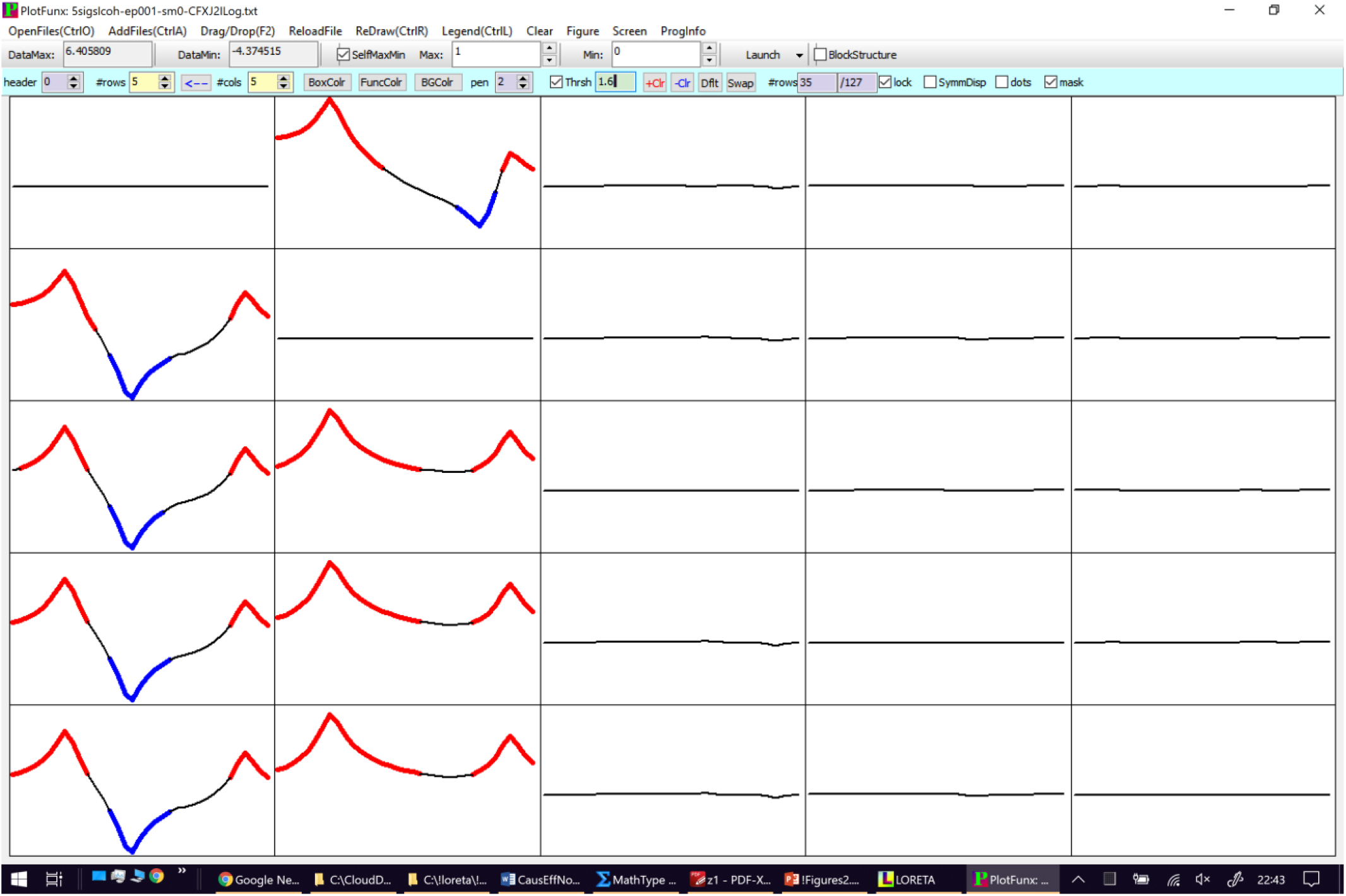
The CFX curves correspond to the difference of green minus magenta curves from Figure 4, i.e. to the difference of log-spectra of the intact system minus the edited system. The curve in the i-th row and j-th column corresponds to the causal effects of the j-th node on the i-th node. The curves were thresholded for a large effect size (see Eq. 13), where red color corresponds to positive values of CFX, and blue color to negative values of CFX. The horizontal frequency axis spans 1 to 35 Hz. The vertical axis spans −4.4 to +6.4

Figure 4 shows a matrix of curves of size 5×5. The magenta colored curve in the i-th row and j-th column corresponds to the power spectra at the i-th node in the edited system where the j-th node has been deleted (obtained from Eq. 4). The green colored curve in the i-th row corresponds to the power spectra at the i-th node in the intact system, obtained from Eq. 3. Note that all green curves for the intact system in the i-th row are identical, independent of the column location. Curves in color black correspond to overlap of the two spectra for the edited and intact systems.

The CFX curves correspond to the difference of green minus magenta curves from Figure 4, i.e. to the difference of log-spectra of the intact system minus the edited system, and are displayed in Figure 5. The curve in the i-th row and j-th column corresponds to the causal effects of the j-th node on the i-th node. The curves were thresholded for a large effect size (see Eq. 13), where red color corresponds to positive values of CFX, and blue color to negative values of CFX.

From column 1 in Figure 5, it can be seen that the role of node 1 in the system consists of activating 8 Hz and 32 Hz oscillations at nodes 2, 3, 4, and 5, and also consists of deactivating 17 Hz oscillations at nodes 2, 3, 4, and 5.

From column 2 in Figure 5, it can be seen that the role of node 2 in the system consists of activating 8 Hz and 32 Hz oscillations at nodes 1, 3, 4, and 5, and also consists of deactivating 17 Hz oscillations at node 1. This was already shown and explained in detail in Figure 3 (C).

From columns 3, 4, and 5 in Figure 5, it can be seen that nodes 3, 4, and 5 do not have causal effects on other nodes of the system.

As can be seen from the connectivity of the system shown in Figure 1, nodes 1 and 2 have very large causal effects on other nodes, while nodes 3, 4, and 5 have none.

## 9. A generalization: causal effects (CFX) by system editing for networks

In this section, a network is defined as a set of nodes. The multivariate time series is now partitioned into “*N*” networks, where each network contains at least one node, and where each node belongs to only one network. The number of nodes in a network need not be the same for all networks.

The question of interest here is: What is the role of one network in a system composed of many interconnected networks?

The answer to this question, based on system editing, requires the deletion of all nodes belonging to one network from the MAR parameters of the intact system, and the re-calculation of the cross-spectral matrix for all remaining networks. This is a straightforward generalization of Eq. 4 above.

Let 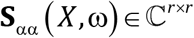 denote the Hermitian cross-spectral matrix for the α-th network in the intact system in Eq. 3, where “r” denotes the number of nodes in the α-th network, with α = 1…*N*.

Let 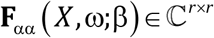 denote the Hermitian cross-spectral matrix for the α-th network in the edited system in Eq. 4, from which all nodes belonging to the β-th network have been deleted, with α ≠ β. In this case, the deletion operator for the parameter matrices sets to zero all rows and columns corresponding to the set of nodes in the β-th network.

Note that **S**_αα_(*X*, ω) and **F**_αα_ (*X*, ω; β) are block diagonal Hermitian matrices contained in Eq. 3 and Eq. 4.

Finally, “causal effects by system editing for networks” is defined as the (causal) change of cross-spectra from the system without the β-th network, to the system with the β-th network. One such measure is:

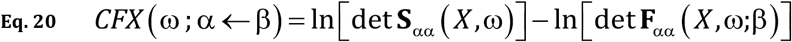

for all networks “α”, such that α ≠ β. In Eq. 20, “det” denotes the determinant of a matrix.

*CFX*(ω; α ← β) in Eq. 20 quantifies the causal effect of network “β” on the observed activity at network “α”.

Note that, as before, this measure can be positive or negative. This means that the addition of a deleted network can causally increase or decrease the determinant of the cross-spectral matrices of the other networks, differently for each frequency.

Also, as before, note that one useful property of the difference of logarithms in Eq. 20 is its invariance to scale, which makes it dimensionless, allowing a comparison of effects on all networks, independent of scale and of network sizes.

## 10. Conclusions, generalizations, final notes

The novel measure proposed here, namely the measure for causal effects by system editing (CFX), offers information on the causal influence that one node (or network) has on the dynamics of the rest of the system.

The measure can be applied to signals other than brain activity.

Nodes or networks can be ranked, in terms of how strongly they affect the system. Such a ranking does not in any way imply that a node with small influence on the system is of little importance.

These measures can in principle be generalized to non-stationary models, and under obvious constraints, to non-linear models.

For the sake of reproducible research, program codes (PASCAL), executable file, and toy data in human readable format are included in the supplementary material.

## Supporting information

PASCAL codes, executable, and toy data (human readable)

