## Supplementary material for "Measuring Granger-causal effects in multivariate time series by system editing": PASCAL codes, executable, and toy data (human readable): CFXhelp.pdf

2018-12-30, Osaka

This refers to the work entitled:
Measuring Granger-causal effects in multivariate time series by system editing

Citation:
Pascual-Marqui, Biscay, Bosch-Bayard, Faber, Kinoshita, Kochi, Milz, Nishida, Yoshimura. Measuring Granger-causal effects in multivariate time series by system editing. 2018-12-21. bioRxiv 504068; doi: https://doi.org/10.1101/504068. [CC-BY-NC-ND 4.0 International license]

The program codes and executable file (CFX.exe) are included.

The program runs under windows.

The programming language is PASCAL.

The compiler is delphi embarcadero community edition version 10.3 called Rio. At the moment of this writing and compilation, it is free for non-commercial stuff.

Run the executable, and make it look like this:

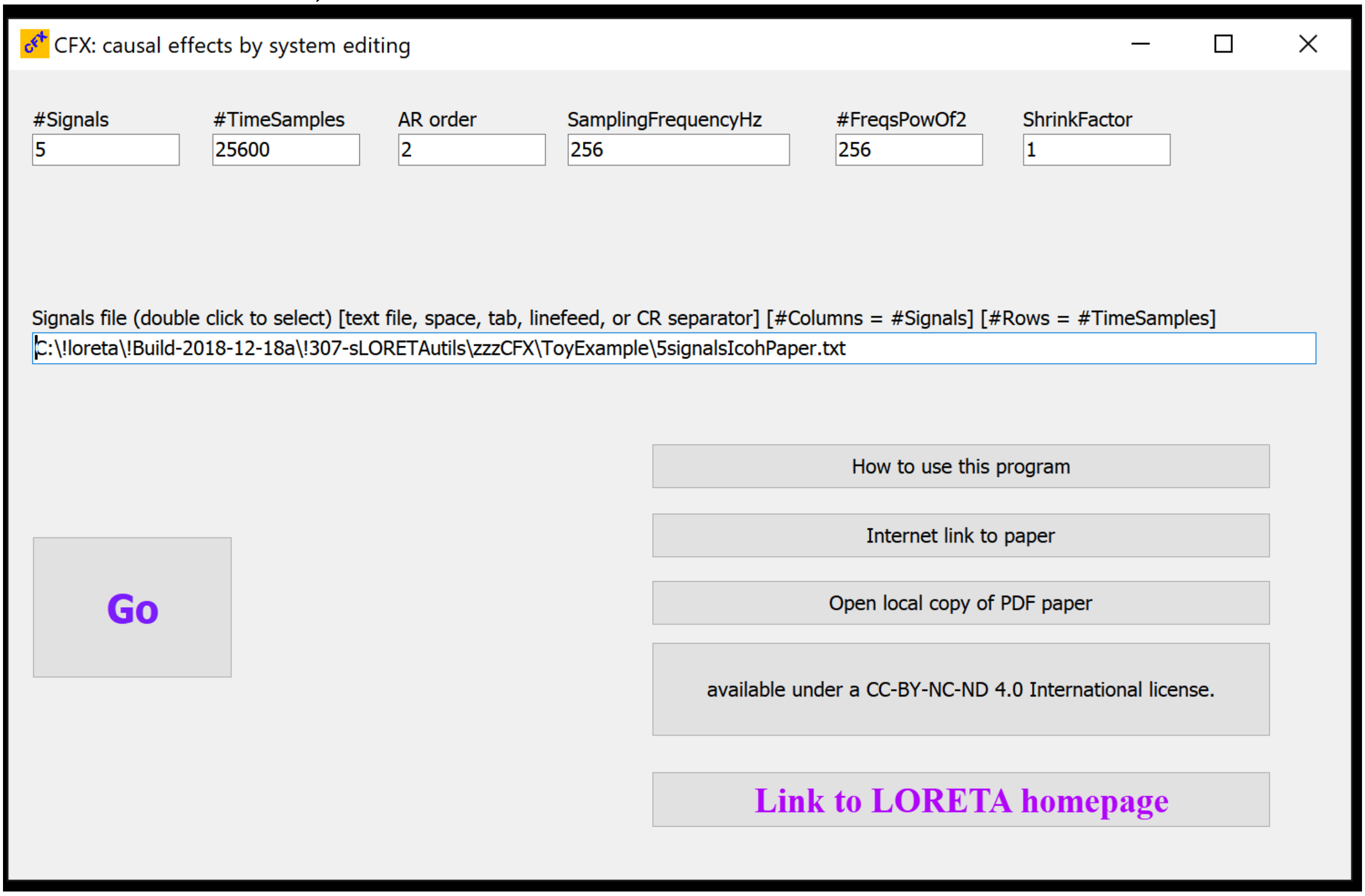


User input:
#Signals: Number of signals.
#TimeSamples: Number of time samples.
AR order: autoregressive order.
SamplingFrequencyHz: Sampling frequency in Hz

#FreqsPowOf2: Number of frequencies, must be a power of 2. The number of discrete frequencies that are prodiced in the output is "(#FreqsPowOf2/2)-1".

ShrinkFactor: Shrink factor must by a number larger than zero and smaller than 1. This is used for regularization, as in "Ridge Regression", for estimating the multivariate autoregressive (MAR) coefficients. ShrinkFactor multiplies the off-diagonal elements of the cross-covariance matrix. ShrinkFactor=1 implies no regularization. ShrinkFactor<1 starts to regularize.

Signals file: double click to select the text file (human readable format) with the signals. The text file should have number of columns equal to the number of signals; and number of rows equal to the number of time samples. The separators between numbers should be space or tab.

The toy example used in the publication corresponds to the signals file named "5signalsIcohPaper.txt", located in the subfolder "ToyExample". This file has 5 dignals, 25600 time samples, and is sampled at 256 Hz. The MAR order is 2.

After you make the program look like the Figure above, click "Go". This produces 2 files, all with the same base filename, but with different extensions. All files are text file, human readable:

5signalsIcohPaper-FreqList.txt: This is the list of the discrete frequencies in Hz of all other output files.

5signalsIcohPaper-CFXJ2ILog.txt: This is the main output file, with the causal effects. It consists of "(#FreqsPowOf2/2)-1" matrices, one for each discrete frequency. and one under the other. Each matrix is of size #Signals*#Signals. For the k-th discrete frequency, the element in the i-th row, j-th column is the effect of the j-th node on the i-th node.

That's it! Everything else is explained in the paper cited above.
